# Gene markers for exon capture and phylogenomics in ray-finned fishes

**DOI:** 10.1101/180786

**Authors:** Jiamei Jiang, Hao Yuan, Xin Zheng, Qian Wang, Ting Kuang, Jingyan Li, Junning Liu, Shuli Song, Weicai Wang, Fangyaun Cheng, Hongjie Li, Junman Huang, Chenhong Li

**Author notes:** Shanghai Universities Key Laboratory of Marine Animal Taxonomy and Evolution Key Laboratory of Exploration and Utilization of Aquatic Genetic Resources (Shanghai Ocean University), Ministry of Education, Shanghai 201306, China National Demonstration Center for Experimental Fisheries Science Education (Shanghai Ocean University). Corresponding author: Chenhong Li.

## Abstract

Gene capture coupled with the next generation sequencing has become one of the favorable methods in subsampling genomes for phylogenomic studies. Many target gene markers have been developed in plants, sharks, frogs, reptiles and others, but few have been reported in the ray-finned fishes. Here, we identified a suite of “single-copy” protein coding sequence (CDS) markers through comparing eight fish genomes, and tested them empirically in 83 species (33 families and 11 orders) of ray-finned fishes. Sorting through the markers according to their completeness and phylogenetic decisiveness in taxa tested resulted in a selection of 4,434 markers, which were proven to be useful in reconstructing phylogenies of the ray-finned fishes at different taxonomic level. We also proposed a strategy of refining baits (probes) design a posteriori based on empirical data. The markers that we have developed may fill a gap in the tool kit of phylogenomic study in vertebrates.

## Introduction

Next-generation sequencing (NGS) drastically reduced the cost of genome sequencing, so that reconstructing phylogenetic relationships using whole genomes became feasible (Jarvis, et al. 2014). However, sequencing whole genomes is still costly and sometime unnecessary. Subsampling genome sequences has gained popularity in phylogenomics and population genomics in recent years (Emerson, et al. 2010; Faircloth, et al. 2012; Lemmon, et al. 2012; Peterson, et al. 2012; Li, et al. 2013). There are two camps that prefer different genome subsampling tools. One is associated with restriction site related markers, such as restriction site associated DNA (RAD) (Baird, et al. 2008) and double digest RADseq (ddRAD) markers (Peterson, et al. 2012), which could be used to produce sequences from a tremendous number of anonymous loci, particularly useful in studying population genomics or species-level phylogeny (Davey and Blaxter 2010). The other camp uses methods of gene capture, also known as target enrichment to capture and sequence target loci, which often result in less missing data than the restriction site related methods does (Collins and Hrbek 2015), and the target loci can be applied across highly divergent taxonomic groups (Faircloth, et al. 2012; Lemmon, et al. 2012; Li, et al. 2013).

Gene capture is based on hybridizing RNA/DNA baits (probes) to DNA library of targeted species and pulling out sequences similar to the baits for subsequent high-throughput sequencing. Two popular methods, Ultraconserved Element Captures (UCE) (Faircloth, et al. 2012) and Anchored Hybrid Enrichment (AHE) (Lemmon, et al. 2012) were developed to pull out highly conserved elements in the genome along with variable flanking regions. Both UCE and AHE methods were designed to anchor highly conserved regions of the genome and make use of variation in flanking sequences. A third method, exon capture was designed explicitly to capture single-copy coding sequences across moderate to highly divergent species (Bi, et al. 2012; Hedtke, et al. 2013; Li, et al. 2013). The advantage of exon capture is that exon sequences are easier to align and better studied for phylogenetics than anonymous non-coding regions. Furthermore, lowered stringency in hybridization and washing steps of exon capture can generate data from more loci than methods focused only on highly conserved elements.

Exon capture markers have been developed in plants (Mandel, et al. 2014; Weitemier, et al. 2014; Chamala, et al. 2015), invertebrates (Hugall, et al. 2016; Mayer, et al. 2016; Teasdale, et al. 2016; Yuan, et al. 2016), and many vertebrate groups, including sharks and skates (Li, et al. 2013), frogs (Hedtke, et al. 2013; Portik, et al. 2016), skink lizards (Bragg, et al. 2016) and others, yet few exon markers have been reported in the ray-finned fishes (Actinopterygii), the most diverse group of vertebrates with more than 30,000 described species (Nelson, et al. 2016). Ilves and Lopez-Fernandez (2014) developed 923 exon markers for cichlids based on genome sequence of tilapia, but those makers probably are too specialized to be used on other ray-finned fishes. We also developed 17,817 single-copy nuclear coding (CDS) markers and applied those in the sinipercid fish, but those markers have not been tested in other ray-finned fishes (Song, et al. 2017).

Selecting target markers and designing baits that are effective across a wide range of species is the first major challenge when applying the gene capture method. Many considerations have been taken into baits design, such as uniqueness and conserveness of markers, length and complexity of markers, and genetic distance between baits and target sequences (Bi, et al. 2012; Faircloth, et al. 2012; Lemmon, et al. 2012; Li, et al. 2013; Mayer, et al. 2016). However, all these measures were taken a priori, and nothing has been done to refine baits design after gene capture to improve the baits set for future experiments.

In this study, we tested the 17,817 CDS markers that we have developed in a previous study (Song, et al. 2017), and screened for the best markers for all major ray-finned fish clades. We chose the best markers according to results of pilot experiments and refined the baits design to improve evenness of reads coverage in different loci. Finally, we tested phylogenetic usefulness of selected markers in ray-finned fishes at both high taxonomic level and species level. Our goal is to provide a set of common exon markers for gene capture and phylogenomic studies in the ray-finned fishes.

## New Approaches

### Testing the targeted gene markers in different groups of ray-finned fishes

We tested the single-copy CDS markers identified from our previous study (Song, et al. 2017). The markers were identified through comparing eight fish genomes (Fig. 1A) using a bioinformatics tool, EvolMarkers (Li, et al. 2012) (supplementary materials Fig. S1). Baits designing steps can be found in detailed materials and methods of supplementary materials. Thousands of the candidate CDS markers were tested empirically in 83 actinopterygian species (99 individuals, 33 families of 11 orders), covering major clades of ray-finned fishes (supplementary materials Table S1). The species captured were part of five different research projects conducted in the authors’ laboratory, including works on basal actinopterygians (Basal), acipenseriforms (Acipen), ostarioclupeomorphs (Ostario), gobioids (Goby) and sinipercids (Sini) (supplementary materials Fig. S2).

**Fig. 1.**
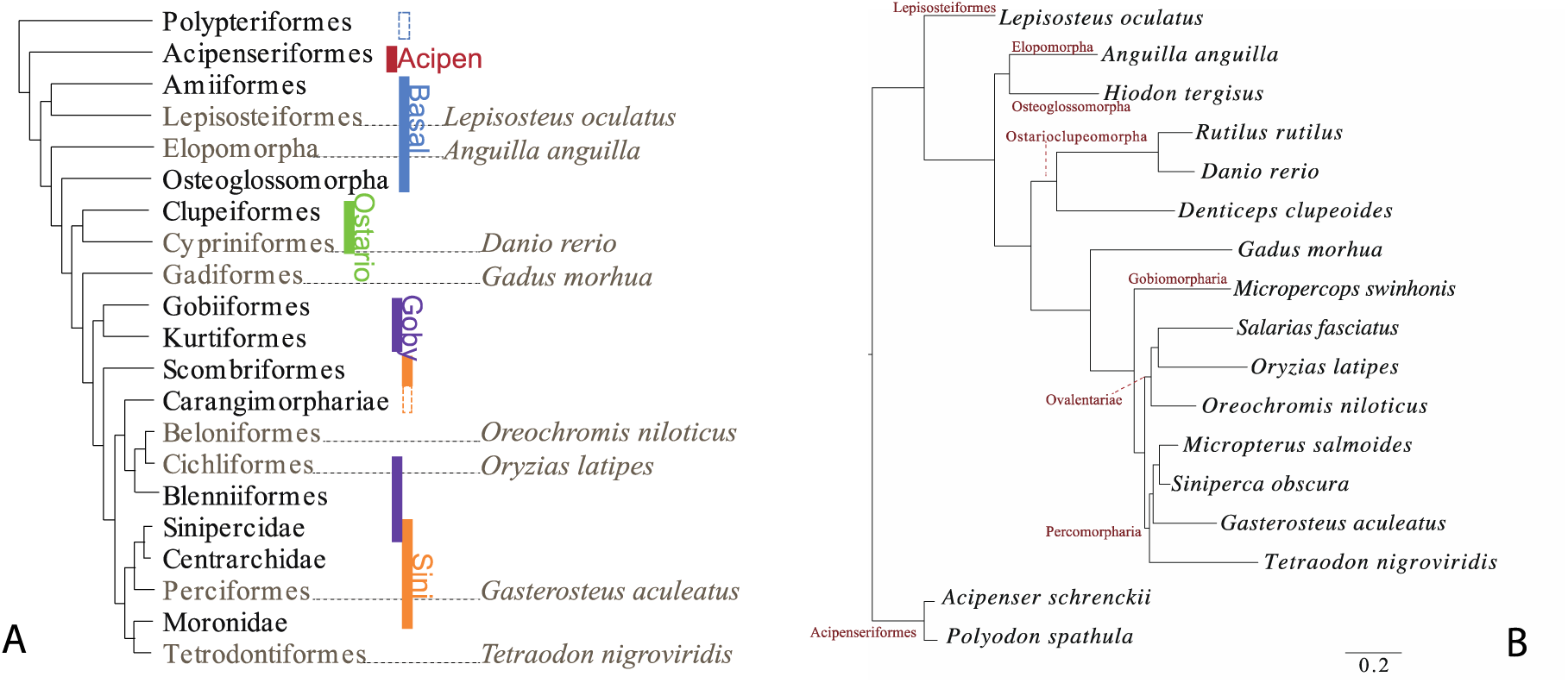
A. Phylogenetic relationships among 21 groups of ray-finned fish (Betancur, et al. 2013). Eight species names indicate the fishes with genome sequence available that were used in finding the target markers. The vertical bars indicate different projects carried in the author’s laboratory. The unfilled vertical bars indicate groups that captured less than 3000 loci. B. Maximum likelihood tree of 16 representative ray-finned fishes based on 4,434 exon loci, all nodes have a 100 bootstrap value.

### Selecting the best markers and refining the baits design based on gene capture results

Based on results of the pilot experiments, target gene markers and baits were evaluated and redesigned to improve their efficacy. There were two major considerations: 1) to select for markers which resulted in less missing data and were phylogenetically decisive, and 2) to identify regions with extraordinarily high read depth and mask those regions for future baits design (Fig. 2). The assembled sequences from different projects were merged (*merge.pl*). Taxa had more than 3,000 genes captured were kept (*select.pl*). Subsequently, a Perl script *deci.pl* was used to pick phylogenetically decisive loci. Phylogenetic decisiveness means that the data sets should contain all taxa whose relationships are addressed (Dell’Ampio, et al. 2014). In our case, the decisive taxonomic groups included eight major clades of the ray-finned fishes: Acipenseriformes, Lepisosteiformes, Elopomorpha, Osteoglossomorpha, Ostarioclupeomorpha, Gobiomorpharia, Ovalentariae and Percomorpharia. The Polypteridae was excluded in bait design, because both species of the polypterids sampled had less than 3000 targets captured.

**Fig. 2.**
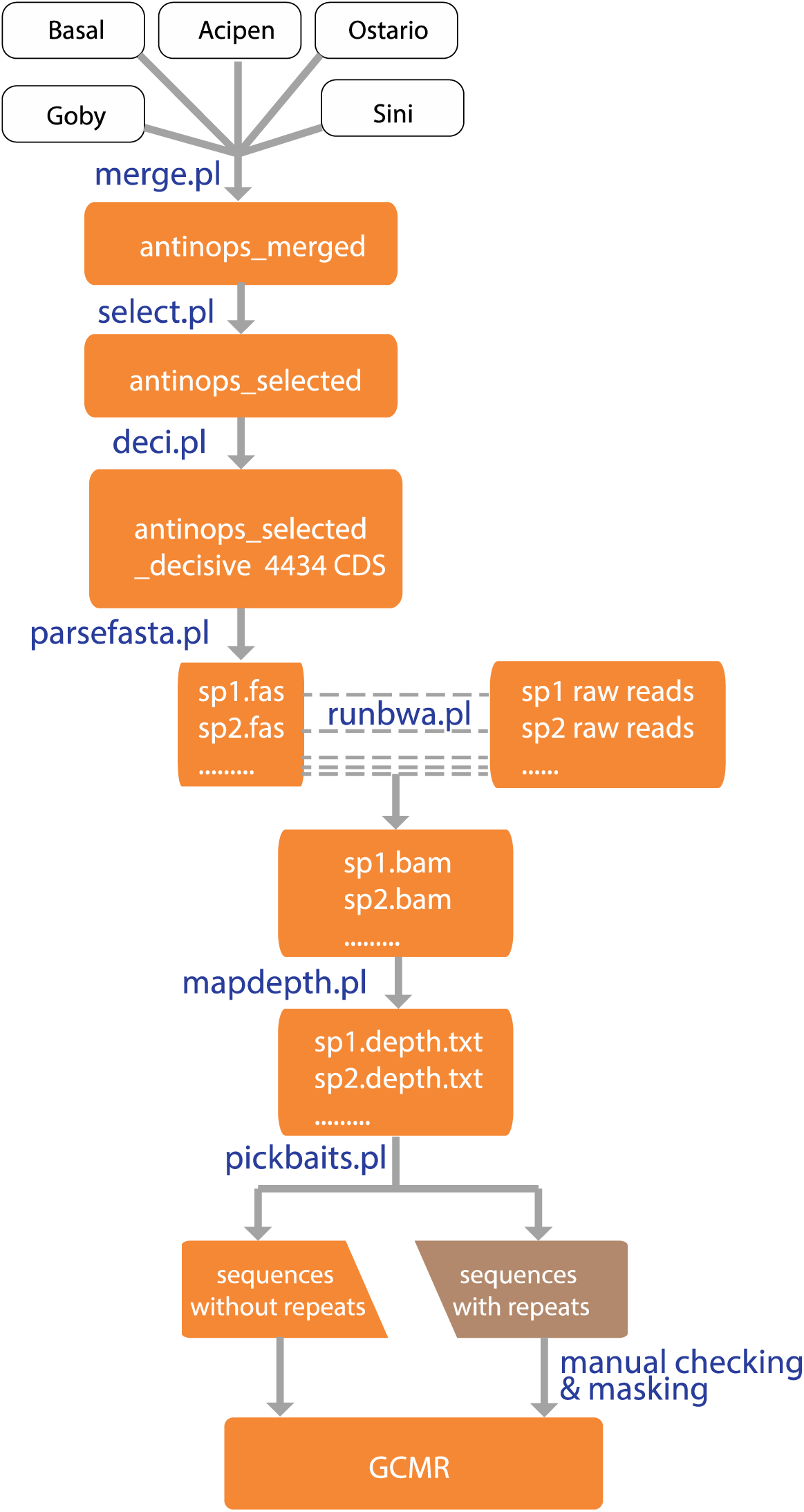
Pipeline of screening for markers with less missing data and better phylogenetic decisiveness and posterior baits refining. I. Merge data from different project (merge.pl); II. select loci with less missing data and high phylogenetic decisiveness (select.pl; deci.pl); III. find and mask region with extraordinary read depth for bait redesign (parsefasta.pl; runbwa.pl; mapdepth.pl; pickbaits.pl). The posterior baits refining steps are optional when empirical data from pilot gene capture are available. GCMR stands for gene capture marker refinement.

From our previous experience, we found that partial regions of some target loci had extraordinarily high number of reads mapped, which consumed a large proportion of the total data collected. Those regions escaped RepeatMasker (Smit, et al. 1996-2004) checking in original baits design, wasted a lot of sequencing reads and are better to be excluded from future baits design. To find those problematic regions, the selected decisive data were parsed to different files by species name (*parsefast.pl*), then, the raw reads of each species were mapped to the assembled reference sequences of each species using *BWA* (Li and Durbin 2009). The reads depth data were extracted from the mapping results using *SAMtools* (Li, et al. 2009) and a custom Perl script (*mapdepth.pl*). Regions with extraordinary high read depth, i.e., 100 times than adjacent regions were identified (*pickbaits.pl*), and manually checked and masked for future baits design. All custom Perl scripts can be found at http://www.lmse.org/markersandtools.html.

### Testing phylogenetic usefulness of the markers selected and efficacy of the new baits

A phylogeny of 16 orders of ray-finned fishes, including 10 species with gene captured data and 7 species with sequence data extracted from genomes were reconstructed. A phylogeny of four species of freshwater sleepers (*Odontobutis*, Gobiiformes) also was reconstructed based on gene capture data of the chosen markers, including five individuals of each species of *Odontobutis sinensis*, *O. potamophila* and *O. yaluensis* and one individual of *O. haifengensis*. Two individuals of *Perccottus glenii* were used as outgroup. Therefore, the phylogenetic usefulness of the chosen markers was evaluated in reconstructing phylogenies of ray-finned fishes at both high and low taxonomic levels. Additionally, we extracted single nucleotide polymorphisms (SNPs) from captured data of the *Odontobutis*, and visualized inter- and intra-specific genetic variation among individuals of the four *Odontobutis* species using the principal component analysis (PCA).

The new baits refined based on empirical data were compared with the baits designed a priori. Reads depth and evenness of reads coverage were summarized from the gene capture data. The comparison was done on results of capturing a goby species (*Rhinogobius giurinus*). Finally, to help researchers to design baits using reference species that are closer to their organism of interested than the eight model fishes that we used, we developed a pipeline of retrieving sequences of the target loci from user provided genomes (supplementary materials file 3).

## Results

### Single-copy protein coding markers for ray-finned fishes

The number of loci captured ranged from 435 to 11,534 in different samples. All but four samples had more than three thousand loci captured (supplementary materials Fig. S2). The samples did the worst in gene capture experiment included two polypteriforms (*Erpetoichthys calabaricus* and *Polypterus endlicher*), one sturgeon (*Acipenser ruthenus*) and the Waigeo barramundi (*Psammoperca waigiensis*). After combining the data from all five projects, excluding taxa with less than 3,000 loci captured and selecting for phylogenetic decisive loci, we obtained 4,434 CDS markers of 2,261genes. The information of the target loci and sequences of the eight model fish species can be found at http://www.lmse.org/markersandtools.html.

### Phylogenetic usefulness of selected markers

The average length of coding region of the chosen markers was 236 bp (94 bp to 4,718bp). GC content ranged from 37% to 69% with an average of 55%. Average pairwise distance (p-dist) among the 17 species varied from 0.06 to 0.50 substitutions per site, with an overall average of 0.19. Average consistency index (CI) was 0.60 (0.43-0.93), and average retention index (RI) was 0.52 (0.47-0.62) (supplementary materials Fig. S3). Maximum likelihood (ML) analyses concatenating 4,434 loci resulted in a well-resolved tree of major ray-finned fish clades, and all nodes had 100 bootstrap support values (Fig. 1). The resulting phylogenetic tree is consistent with recent studies (Betancur, et al. 2013; Faircloth, et al. 2013), except that the Elopomorpha and the Osteoglossomorpha were found sister to each other.

There were 4,296 of 4,434 loci captured at least in one *Odontobutis* sample. A total of 1,630 loci were captured in all samples. The average length of target regions was 265 bp (120 bp to 5,637 bp). The average length of captured non-coding flank region was 487 bp. A concatenated ML tree was reconstructed for the four Chinese *Odontobutis* species with *P. glenii* as the outgroup. The species level phylogeny was well resolved with 100 bootstrap support values for each node. *Odontobutis sinensis* was sister to the rest of *Odontobutis* species. *Odontobutis yaluensis* was grouped with *O. potamophila* and *O. haifengensis* was placed as sister to them. Species tree is consistent with ML tree with a normalized quartet score 0.64 (supplementary materials Fig. S4). We extracted 36,440 single nucleotide polymorphisms (SNPs) sites from target regions (35 SNPs per kb). In PCA, axis 1 and axis 2 explained 48.42% and 11.21% of the variability respectively. Individuals of *O. yaluensis* and *O. potamophila* were close to each other, whereas individuals of *O. sinensis* were apart from them and *O. haifengensis* lied in between (supplementary materials Fig S5).

### Gene-capture marker refinement

We examined the results of gene capture experiments using original baits. We found that 26 loci of *Rhinogobius giurinus* had extreme high number of reads mapped. We manually checked those loci and found that all regions with high reads depth had low complexity. We masked those regions, redesigned the baits and carried a new round of gene capture experiment. The gene capture results from new baits had better even coverage among different loci than the results from the original baits (Fig. 3).

**Fig. 3.**
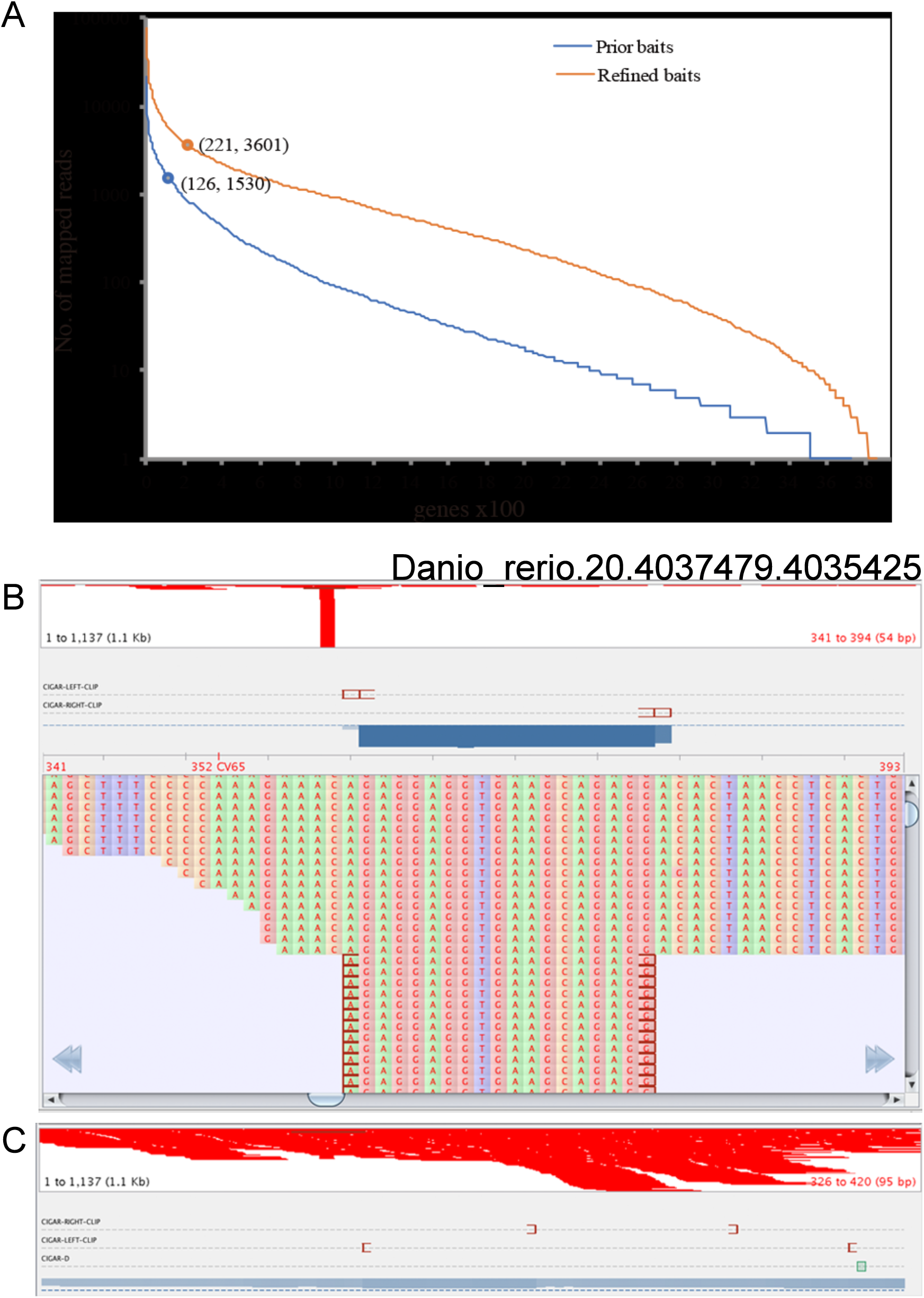
Comparison on evenness of read coverage between results of gene capture using the baits designed a priori (A, blue curve) and the baits refined posteriorly (A, orange curve). B and C are screenshots from visualizing the read depth of the locus Danio_rerio.20.4037479.4035425 using Tablet v1.16.09.06. In this example, the result using baits designed a priori (B) is much worse than the result using refined baits (C).

## Discussions

### Exon capture

Protein-coding sequences are easy to align and molecular evolution of protein sequence is better studied than non-coding flank regions, whose variation tend to increase when further apart from the conserved core region (Faircloth, et al. 2012). Our experiments showed that the markers selected and the baits designed were effective in studying phylogenetic relationship of major groups of the ray-finned fishes, and closely related species as well. We notice that our exon capture protocol also produced data from flanking non-coding regions with an average length of 487 bp. We did not analyze the sequence data from the flanking region, because the non-coding flanking regions of many loci could not be aligned. Further investigation on how to process and utilize the data of flanking regions for studies at inter- and intraspecific level should be carried out.

### A posteriori marker design

The simple repeats in the markers were detected and masked using RepeatMasker by the manufacturer, MYcroarray (Ann Arbor, Michigan) before synthesizing the baits. However, repeats with some variations or complex repeats could not be detected with RepeatMasker, thus resulted in a high read depth in some regions (Fig. 3B). Extreme high read depth suggests that many reads were not from the target regions, which could cause problem in subsequent read assembly and waste sequencing resource. Based on the sequencing results, we masked these unusual regions in the following baits refinement in gobies, which has shown more even coverage for the targeted loci (Fig. 3B). If a pilot study is planned before a large-scale experiment, we recommend applying our method to refine baits design to improve the efficacy of baits.

### Orthology checking and data filtering

Problem of mistakenly using paralogous genes for phylogenetic reconstruction is exacerbated with phylogenomic data, and currently there is no ideal method to validate orthology of loci assembled from NGS data (McCormack, et al. 2013; Chakrabarty, et al. 2017). The targeted loci we selected for are “single-copy” (Li, et al. 2012), which may have less chance to be paralogous than members of gene families, (Li, et al. 2007). In addition, we performed a “re-blast” step in data processing pipeline to identify and exclude potential paralogs (Yuan, et al. 2016). Nonetheless, both method cannot guarantee orthology of targeted sequences due to the third round of whole-genome duplication event in teleost and slow and steady loss of some paired genes in the subsequent 250 My (Inoue, et al. 2015). Tree based methods, such as filtering the loci a posteriori based on known monophyly of taxa could be used to alleviate the problem of paralogy.

## Materials and Methods

For detailed materials and methods, see supplementary materials file 4.

## Supplementary Materials

Supplementary File 1: Figures S1 - S5.

Supplementary File 2: Tables S1.

Supplementary File 3: A user-friendly pipeline to retrieve target sequences of the 4,434 loci from new genome sequences or transcriptomes.

Supplementary File 4: Detailed materials and methods.

## Acknowledgements

This work was supported by the Innovation Program of Shanghai Municipal Education Commission and the Program for Professor of Special Appointment (Eastern Scholar) at Shanghai Institutions of Higher Learning. We would like to thank Shanghai Oceanus Supercomputing Center (SOSC) for providing computational resource.

